# Genetically-encoded markers for confocal visualization of single dense core vesicles

**DOI:** 10.1101/2024.10.07.617131

**Authors:** Junwei Yu, Yunpeng Zhang, Kelsey Clements, Nannan Chen, Leslie C. Griffith

## Abstract

Neuronal dense core vesicles (DCVs) store and release a diverse array of neuromodulators, trophic factors and bioamines. The analysis of single DCVs has largely been possible only using electron microscopy, which makes understanding cargo segregation and DCV heterogeneity difficult. To address these limitations, we developed genetically-encoded markers for DCVs that can be used in combination with standard immunohistochemistry and expansion microscopy, to enable single-vesicle resolution with confocal microscopy.

## Main

Release of neuroactive substances in the brain has classically been thought to occur via two distinct pathways. Small-molecule neurotransmitters, packaged into small clear synaptic vesicles (SVs, 30-40 nm diameter), are released at active zones of synapses. In contrast, peptide and neuromodulators are packaged into dense core vesicles (80-200 nm diameter) which fuse extrasynaptically^1^. Neuromodulators play crucial roles in transducing the effects of internal states and external conditions to the brain, making understanding the mechanisms of neuromodulator release essential for understanding how context influences behavior^2^.

Co-transmission, the release of multiple neuroactive substances by single cells, introduces another level of complexity. Co-packaging of multiple substances into a single vesicle imposes different constraints on signaling compared to the situation in which a cell can traffic and release each substance independently. Understanding where a neurochemical is released and what other substances are co-released is crucial for comprehending the interactions between synaptic and modulatory pathways. These questions have most often been addressed using techniques with single-vesicle resolution, e.g. single synapse functional data^3^ or immuno-electron microscopy^4^. While these techniques can observe specific synapses, they do not allow for a comprehensive examination of the occurrence of co-packaging and co-transmission.

Here we develop genetic tools for DCVs visualization, enabling single DCV resolution with light microscopy when combined with expansion microscopy (ExM)^5^. IA2 family proteins (PTPRN and PTPRN2 in mammals, IDA1 in *C. elegans,* IA2 in *Drosophila*) are trans-membrane proteins of DCV that are expressed in neuroendocrine cells throughout the body, making them excellent markers for DCVs^6^. Consistent with this, CRISPR/Cas9 insertion of monomeric green fluorescent protein (mEGFP) into the *Drosophila IA2* genetic locus to produce a C-terminus fusion (Extended Data Fig. 1a), demonstrated widespread expression of IA2 in both adult and larval brains (Extended Data Fig. 1b).

We utilized the GAL4/UAS system^7^ for cell-specific expression (Fig. 1a). *Pigment-Dispersing Factor (PDF)-GAL4,* a driver expressed in peptidergic ventrolateral neurons (LNvs) of the *Drosophila* circadian clock, demonstrated colocalization of EGFP with PDF peptide (Extended Data Fig. 2a). ExM, which increases brain size by about 4.5-fold, and DCVs to about 360-900 nm in diameter, made DCVs visible with light microscopy (Fig. 1f). We found PDF peptide located at the center of IA2-containing circular structures (Extended Data Fig. 2b-c), suggesting that IA2::mEGFP localizes to DCVs which store and release PDF. Notably, in the small LNv projections we did not observe PDF puncta that lacked adjacent IA2 staining. This is the first time that single dense-core vesicles have been visualized by optical microscopy in tissue.

**Fig 1.**
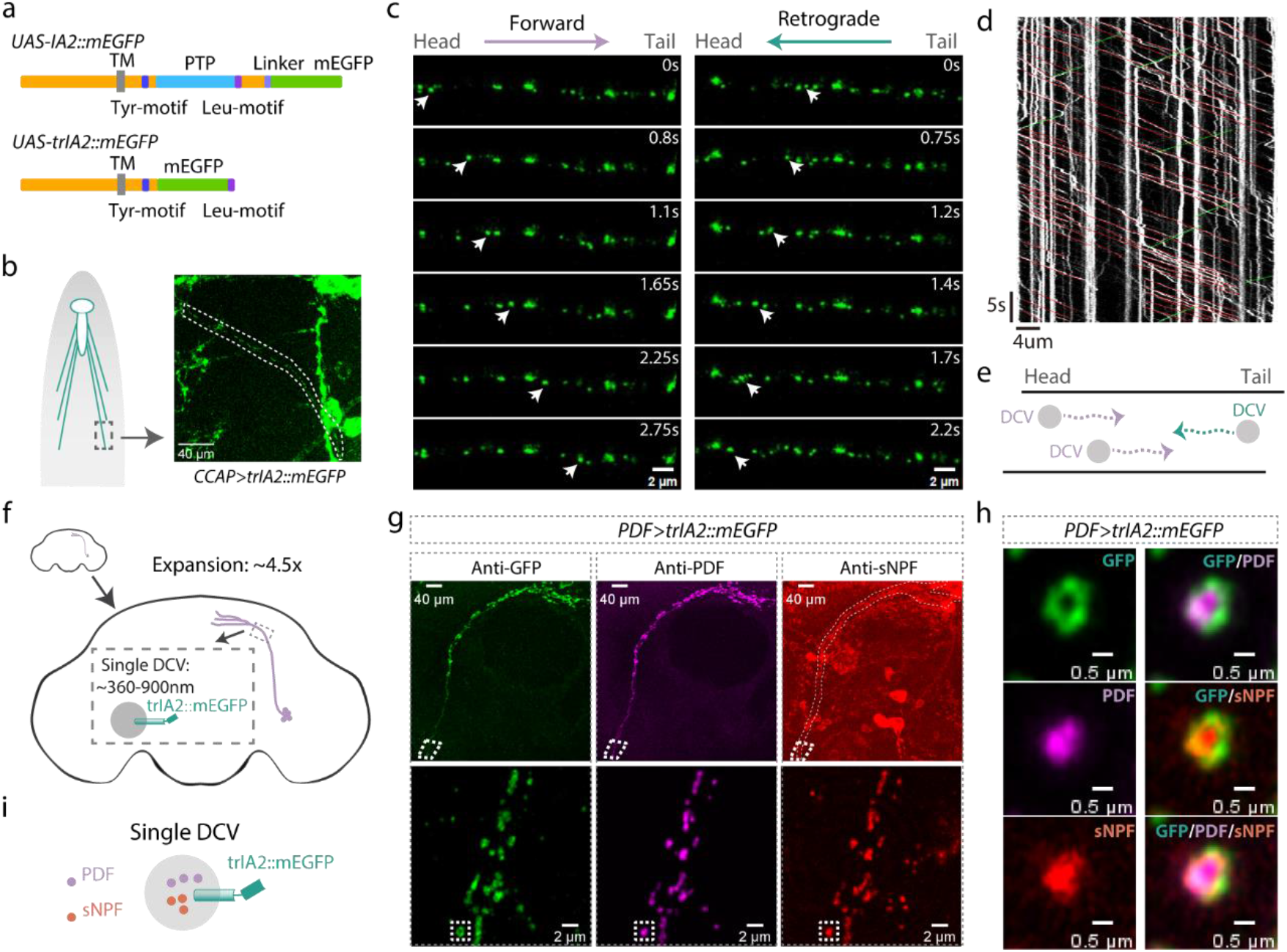
Visualizing individual DCVs. **a**, Schematic diagrams of *Drosophila IA2* transgenes: In the *UAS-IA2::mEGFP* fly, mEGFP is fused to the C-terminus of IA2 (upper panel). In the *UAS-trIA2::mEGFP* fly, the C-terminal PTP domain is removed and replaced with mEGFP, followed by IA2’s Leu-motif (lower panel). TM: transmembrane domain, PTP: protein-tyrosine phosphatase domain. **b**, Cartoon and representative image showing projection (dotted lines) of a trIA2::mEGFP-expressing CCAP neuron. **c**, Sequential images showing vesicles (arrowheads) moving from head to tail (left panels) or tail to head (right panels). Scale bar: 2 µm in each panel. **d**, Image depicts vesicle movement along the motor neuron projection over time. Red: individual vesicles moving forward, green: vesicles moving retrogradely, white: stationary vesicles. **e**, Cartoon illustrating relative levels of vesicle movement. **f**, Cartoon illustrating the approximately 4.5-fold brain size increase, with 360-900 nm DCVs. **g-h**, PDF and sNPF peptides are co-packaged into the same DCVs. Lower panels of **g** show enlarged images of outlined area in upper panels. **h** shows close-up of the inset in lower panels of **g**.Green: mEGFP, magenta: PDF, red: sNPF in **g**-**h**. Scale bar: 40 µm in upper panels of **g**, 2 µm in lower panels of **g** and 0.5 µm in **h. i**, cartoon of co-packaging.

To check whether GAL4-driven expression of IA2::mEGFP affects DCV function, we quantified PDF staining in the projection regions of small LNvs and found there was an increase in PDF signal (Extended Data Fig. 3). This indicates that IA2::mEGFP is functional, but suggests that overexpression of wildtype IA2 increases steady-state DCV levels. Since the protein-tyrosine phosphatase (PTP) region of IA2 is conserved and functionally important, we constructed *UAS-truncated(tr)IA2::mEGFP* lines lacking that domain (Fig. 1a). Expression of trIA2::mEGFP also labeled single DCVs after expansion (Fig. 1g), but did not change PDF levels (Extended Data Fig. 3), making trIA2::mEGFP a better GAL4-driven DCV marker.

We noticed that nearly all IA2 signal visible in small LNv processes has corresponding PDF staining (Fig. 1g and Extended Data Fig. 2), suggesting that IA2 exclusively labels DCVs and not SVs, much like mammalian PTPRN, which is excluded from SVs^8^. To rule out association between fly IA2 and SVs, we generated an IA2 knock out strain by deleting the last eight exons of the *IA2* gene. This line was homozygous viable, and adult brains had a dramatic decrease in DCV cargo-positive puncta in small LNv projections, indicating that IA2 enhances, but is not required for, DCV function. Importantly, the levels of synaptophysin-labeled SVs in LNvs remained unchanged, confirming that IA2 exclusively affects DCVs (Extended Data Fig. 4a-c). Consistently, immunohistochemical localization of trIA2::mEGFP in motor neuron terminals at the larval neuromuscular junction demonstrates that it does not co-localize with cysteine string protein (CSP), an SV marker (Extended Data Fig. 5). Cell-specific loss of IA2 indicates that its role in DCV function is cell autonomous (Extended Data Fig. 4d).

In the cytoplasm of neurons, DCVs are dynamic. To determine if IA2 could be used as a marker in live imaging, we examined the projections of larval motor neurons expressing trIA2::mEGFP (Fig. 1b). We observed labelled DCVs moving from soma to synaptic regions, as well as a few DCVs moving retrograde (Fig. 1c-e). These results indicate that these genetic reagents can also be used to investigate the mechanisms underlying DCV movement in real-time.

Many neurons, including LNvs^9^, express multiple peptides. To determine if our marker could be used to distinguish between co-release from the same DCV and co-transmission via independent DCV populations, we stained adult brains from *PDF>trIA2::mEGFP* animals with antibodies to PDF and sNPF. We found that the peptides locate together at the center of single vesicles (Fig. 1g-i). We found a similar situation in the motor neuron of muscle 12 in the third instar larva (Fig. 1b), where CCAP and pBurs co-localize in the same DCVs (Extended Data Fig. 6). These results demonstrate that multiple neuropeptides can be co-packaged into the same DCVs for co-release in both larval and adult *Drosophila* neurons and that IA2 marker transgenes can be used to distinguish between co-release and co-transmission via multiple DCV pools.

Most well-described DCV cargoes are proteinaceous; small molecules involved in fast neuronal communication are primarily released from SVs. Bioamines are an exception to this rule and are known to be packaged in both SVs and DCVs, reflective of their dual roles as synaptic transmitters and extrasynaptic modulators^10,11^. We wondered whether other small molecule neurotransmitters might also have roles as modulators and be packaged into DCVs. To test this idea, we examined co-localization of IA2::mEGFP with vesicular transporters, proteins localized to vesicle membrane which package neurotransmitters into SVs. Each of the main small molecule neurotransmitters requires a different transporter: vesicular monoamine transporter (VMAT) for bioamines, vesicular acetylcholine transporter (VAChT) for acetylcholine, vesicular glutamate transporter (VGluT) for glutamate and vesicular GABA transporter (VGAT) for γ-aminobutyric acid (GABA). To determine if IA2 was normally present in neurons that release these transmitters, we constructed an *IA2-Frt-stop-Frt-mEGFP* fly strain (Fig. 2a) by inserting an Frt-stop-Frt-mEGFP cassette at the C-terminus of the *IA2* locus. The stop cassette suppresses EGFP tagging unless removed by recombination. Expression of flippase (Flp) cell-specifically fuses the endogenous IA2 protein in *GAL4>Flp* cells with EGFP. We found high levels of endogenous IA2 expression in bioaminergic (*VMAT>Flp*, Fig. 2b), GABAergic (*VGAT>Flp*, Fig. 2c), cholinergic (*VAChT>Flp*, Extended Data Fig. 7a) and glutamatergic (*VGluT>Flp*, Extended Data Fig. 7b) cells.

**Fig 2.**
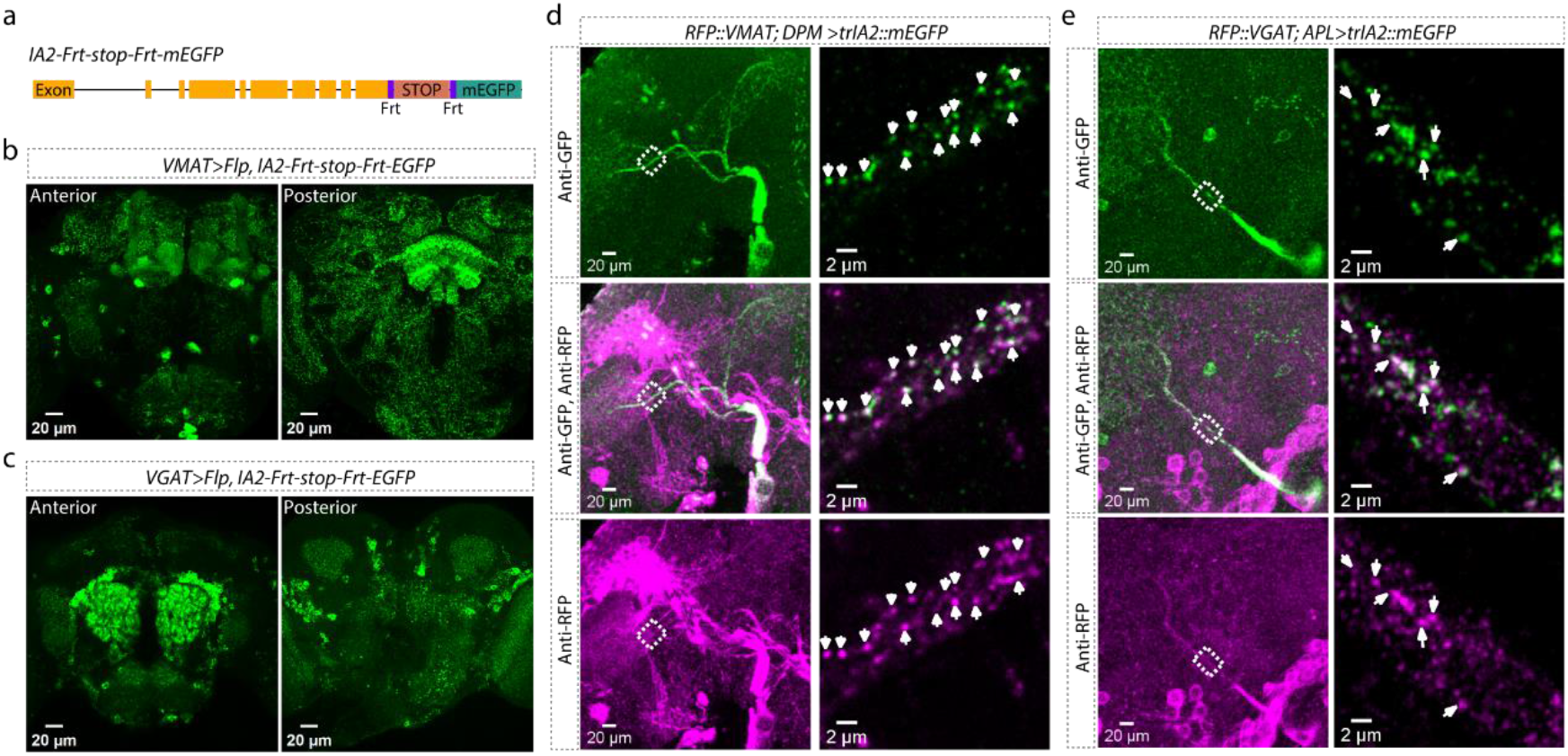
Co-localization of DCV IA2 with VMAT and VGAT. **a**, Schematic showing CRISPR insertion of *Frt-stop-Frt-mEGFP* in the 3’ end of the *IA2* gene. **b-c**, IA2 expression in VMAT-positive (**b**) and VGAT-positive (**c**) neurons. Left panels show anterior view, right panels show posterior view. Scale bar: 20 µm. **d**, Co-localization of RFP::VMAT from endogenous *VMAT* locus with trIA2::EGFP. Left: DPM neuron projections in an expanded fly brain. Right: super-resolution images of the outlined area. Arrowheads indicate DCVs co-labeled by trIA2::mEGFP and RFP::VMAT. Scale bar: 20 µm on left, 2 µm on right. **e**, Co-localization of RFP::VGAT ^<sup>1</sup>^ with trIA2::EGFP. Left: APL neuron projections in an expanded fly brain. Right: super-resolution images of the outlined area. Arrowheads indicate DCVs co-labeled by trIA2::mEGFP and RFP::VGAT. Scale bar: 20 µm on left, 2 µm on right.

Since we knew that VMAT was present in some DCVs^11^, we examined its co-localization with IA2::mEGFP as a positive control. We first used CRISPR/Cas9 to label endogenous VMAT with RFP (*RFP::VMAT*). We then labelled DCVs with trIA2::mEGFP only in the two bioaminergic dorsal paired medial (DPM) neurons^12^. We found that a substantial number of the trIA2::mEGFP puncta also contained VMAT::RFP, confirming the previous biochemical finding that monoamines can be packaged into DCVs (Fig. 2d and Extended Data Fig. 8a).

GABA is known to be present in DCVs in mammalian adrenal^13^. To determine if GABA can be packaged into DCVs in the *Drosophila* brain, we used trIA2::mEGFP to label the DCVs in the GABAergic anterior paired lateral (APL) neurons^14^ on a *VGAT::RFP*^15^ background. Though most VGAT::RFP did not colocalize with trIA2::mEGFP, there were clear instances of co-localized signal, indicating that GABA can be packaged into DCVs (Fig. 2e and Extended Data Fig. 8b). The idea that GABA could be neuromodulatory has been around for a while, and it is clear that extrasynaptic signaling by GABA is important for setting circuit tone in both insect and mammalian brains^13,16,17^. These data suggest that DCVs are a potential source of this modulatory GABA.

While synaptic release of small “fast” transmitters like glutamate, acetylcholine and GABA is relatively well characterized, extrasynaptic release of DCVs has been more difficult to study due in part to the greater diversity of vesicle cargos (peptides, bioamines) and the more subtle circuit functions of neuromodulation. Using the genetic tools we developed, researchers can visualize DCVs, tracking moving DCVs, and observe the co-existence of SVs with DCVs under light microscopy. These tools enable the exploration of a wide variety of questions about the localization and interactions of neurochemical signaling pathways at the whole brain or circuit level.

## Acknowledgments

This work was supported by R21NS096414 and R01NS122970 to LCG. Stocks obtained from the Bloomington Drosophila Stock Center (NIH P40OD018537) were used in this study.

## Author contributions

JY, YZ and LCG designed the experiments. JY, YZ, NC and KC generated reagents and carried out experiments. JY, YZ and NC analyzed data. LCG and NC wrote and edited the manuscript.

## Methods

### Fly strains and husbandry

All flies were raised on standard cornmeal medium at 25°C with a 12h/12h light cycle. For adult fly experiments, flies were collected at eclosion and aged to 3-5 days before performing experiments. *PDF-GAL4* was kindly provided by Dr. Michael Rosbash, *UAS-ANF::mOrange2* by Dr. Edwin S Levitan and *UAS-synaptophysin::pHTomato* by Dr. Andre Fiala. *CCAP-GAL4* (#25685), *ChAT-GAL4* (#60317), *VMAT-GAL4* (#66806), *VGluT-GAL4* (#60312), *nos-GAL4* (#64277) and *UAS-Flp* (#4539) were obtained from Bloomington *Drosophila* Stock Center. *APL-GAL4* (*VT-043924-GAL4*) and *DPM-GAL4* (*VT-064246-GAL4*) were collected from Vienna Drosophila Resource Center. *VGAT-GAL4* and *RFP::VGAT* were constructed in this lab and described previously^1^.

### Generation of 10x*UAS-IA2::mEGFP* and 10x*UAS-trIA2::mEGFP* lines

For the *UAS-IA2::mEGFP* fly strain, the IA2 coding region was amplified from a *Canton-S* wildtype fly cDNA library, and GFP was amplified from pJFRC2-10XUAS-IVS-mCD8::GFP plasmids (Addgene Plasmid #26214) and then amino acid A206 was mutated to K to make mEGFP (monomeric enhanced GFP). For the *UAS-trIA2::mEGFP* fly strain, the PTP domain and the following fragments of *IA2* were deleted and replaced with mEGFP, followed by the Leu-motif. These fragments were assembled in order and subcloned into the pJFRC2-10XUAS-IVS-mCD8::GFP plasmid using the Gibson assembly method (10xUAS-IA2::mEGFP plasmid and 10xUAS-trIA2::mEGFP plasmid in data S1 separately).

These plasmids were verified by sequencing and then injected into *phiC31-attP* flies (Bloomington *Drosophila* stock center, #25710), which have an attP site on the third chromosome to allow targeted integration. The progeny of the injected flies was screened using the w^+^ red eye marker and confirmed by polymerase chain reaction (PCR) and sequencing.

Generation of *IA2-Frt-stop-Frt-mEGFP, RFP::VMAT* and *IA2::mEGFP*

To knock in the *Frt-stop-Frt-mEGFP cassette* at the C-terminus of *IA2*, we designed a guide RNA that recognize the endpoint of *IA2* with an online tool (http://targetfinder.flycrispr.neuro.brown.edu/). This guide RNA was cloned into a pU6 plasmid (Addgene, #45946). Additionally, a donor plasmid (pMC10-IA2-Frt-stop-Frt-mEGFP plasmid in data S1) was created and injected into the Cas9 flies (*y,sc,v; nos-Cas9/CyO; +/+*) along with the gRNA plasmid. Using the same strategy, we knocked in RFP at the N-terminus of VMAT. The guide RNA sequence is listed in table S1, and the donor plasmid is shown in data S1.

To get the *IA2::mEGFP* fly strain, we bred *IA2-Frt-stop-Frt-mEGFP* flies with a stable fly line that constantly expresses Flp from the X chromosome. To get the Flp expressing stable line, we crossed *nos-GAL4* (#64277) with *UAS-FLP* (#29731) flies and obtained one recombinant line. We screened progeny of *nos-GAL4, UAS-Flp;; IA2-Frt-stop-Frt-mEGFP* flies and harvested *IA2::mEGFP* fly strains, in which the Frt sequence was used as a soft linker by adding two nucleotides to the beginning of the first Frt site to make it in frame. For all the lines described above, correct integrations were confirmed by PCR and sequencing using primers that bind outside the integrated junction region.

### Generation of *Frt-IA2::mEGFP-Frt* and *IA2 Null* lines

To generate the *Frt-IA2::mEGFP-Frt* fly strain, we used CRISPR/Cas9 to knock in two Frt sites: one in the third intron of the *IA2* gene and another at the end of *IA2*. Two guide RNAs were designed accordingly and cloned into pU6 plasmids (Addgene, #45946). In the donor plasmid (pMC10-Frt-IA2::mEGFP-Frt plasmid in data S1), mEGFP is inserted at the C terminus of IA2, and followed by the second Frt site. The donor plasmid was co-injected into Cas9 flies (*y,sc,v; nos-Cas9/CyO; +/+*) along with the gRNA plasmids. After obtaining the *Frt-IA2::mEGFP-Frt* fly, we crossed it with *nos-GAL4, UAS-FLP* flies, screened the progeny, and successfully harvested a *IA2 Null* mutant fly strain. This line is homozygous viable. Correct integrations were confirmed by PCR and sequencing using primers that bind outside the integrated junction region.

### Immunohistochemistry and image processing

To dissect and stain the brains of adult and larval flies, we followed the protocols from Janelia (www.janelia.org/project-team/flylight/protocols). Briefly, the brains were dissected in S2 solution and then fixed in 2% paraformaldehyde solution for 55 minutes at room temperature (RT). The brains were then washed four times, 10 minutes each time, with 0.5% phosphate-buffered saline containing Triton X-100 (PBST). Following the washes, the brains were blocked with 5% goat serum in PBST solution for 1.5 hours at RT. The samples were then incubated in primary antibody solution for 4 hours at RT with continued incubation at 4°C over 2-3 nights. Subsequently, samples were washed three times for 30 mins each with 0.5% PBST and incubated in secondary antibody over two nights. The same washing process was performed afterward. Some samples then underwent the expansion protocol as described below, while others are fixed in 4% PFA for an additional 4 hours at RT and mounted in Vectashield mounting medium (Vector Laboratories).

To visualize NMJs on larval body walls, wandering third instar larvae were dissected in cold HL3.1 solution (NaCl 70mM, KCl 5mM, CaCl_2_ 0.1mM, MgCl_2_ 20 mM, NaHCO_3_ 10mM, Trehalose 5mM, Sucrose 115mM, HEPES 5Mm; osmolarity: 395.4 mOsm, pH7.1-7.2) and then fixed in 4% PFA for 10 mins at RT. The samples were then washed in PBST for 3×10 minutes and incubated in primary antibody solution overnight. Following this, the samples were washed again and incubated in secondary antibody solution for another night. After a final wash for 3×30 mins, the mounting process was performed.

The primary antibodies used were rabbit anti-GFP (1:1000; Thermo Fisher Scientific), rabbit anti-RFP (1:200; Takara), mouse anti-GFP (1:200; Sigma-Aldrich), chicken anti-GFP (1:500; Invitrogen), mouse anti-PDF (1:200; Developmental Studies Hybridoma Bank; PDF C7-c), mouse anti-Csp antibody (1:100; Developmental Studies Hybridoma Bank), rabbit anti-CCAP (1:500;Jena Bioscience; ABD-033), rabbit anti-sNPF (1: 500; a gift from Dr. Jan Veenstra, Universite de Bordeaux, France), and mouse anti-pBurs (1:500; a gift from Benjamin White, National institute of Health; originally from Dr. Aaron Hsueh, Stanford University). The secondary antibodies used were Alexa Fluor 488 anti-chicken antibody, Alexa Fluor 488 anti-mouse/rabbit antibody (Invitrogen), Alexa Fluor 561 anti-rabbit and Alexa Fluor 635 anti-mouse/rabbit antibody (Invitrogen), all at 1:200 dilutions. For NMJs staining, Alexa Fluor 488-conjugated anti-GFP antibody (1:250; Invitrogen) was used.

Images were captured using a Leica SP5 confocal microscope with either a 20x or 60x objective lens, except for the NMJs images, which were acquired on a Zeiss LSM880 Airy Scan Fast Confocal System using a 63x objective lens. The images from Leica SP5 were then processed and analyzed using ImageJ Fiji software^2^, while the Airy Scan images underwent deconvolution using Huygens software.

### ExM sample preparation

The brain samples for expansion microscopy were prepared as previously described^3^. After dissecting and staining the brains, they were incubated in AcX solution (0.1mg/ml) for more than 24 hours at RT in the dark. Brains were then washed three times with PBS solution and incubated in gelling solution for 45 minutes on ice in the dark. Gel chambers were constructed by placing two strips of tape approximately 3-4 cm apart on a glass slide. Brains were placed into the gel chambers and incubated in gelling solution at 37°C for 2 hours. After incubation, the brains were trimmed away from the gelling solution and submerged in digestion buffer for 24 hours at room temperature in the dark. Finally, brains were washed with an excess volume of ddH_2_O at room temperature more than three times, 20 mins each time. The samples were then prepared for imaging with a ZEISS LSM 880 Airyscan microscope with a 63x objective.

### Live imaging of DCVs and data analysis

Third instar larval brains were dissected in ice-cold HL3 medium. The brains were then transferred to an imaging chamber containing fresh HL3 saline, which was continuously supplied to the chamber during the recording process. Images of motor neuron projections were captured at 12 Hz with a 63X Multi-Immersion lens under ZEISS LSM 880 Airyscan microscope with the AiryScan FAST model. For the analysis of dense core vesicle trafficking, we used Kymograph plugin in the imageJ^2^ (Fiji) as described previously^4^.

### Statistical analysis

Prism 9 software was used for statistical analysis. Data were tested for normality and then analyzed with either a parametric or non-parametric test as appropriate.

---

**Extended Data Figure 1:**
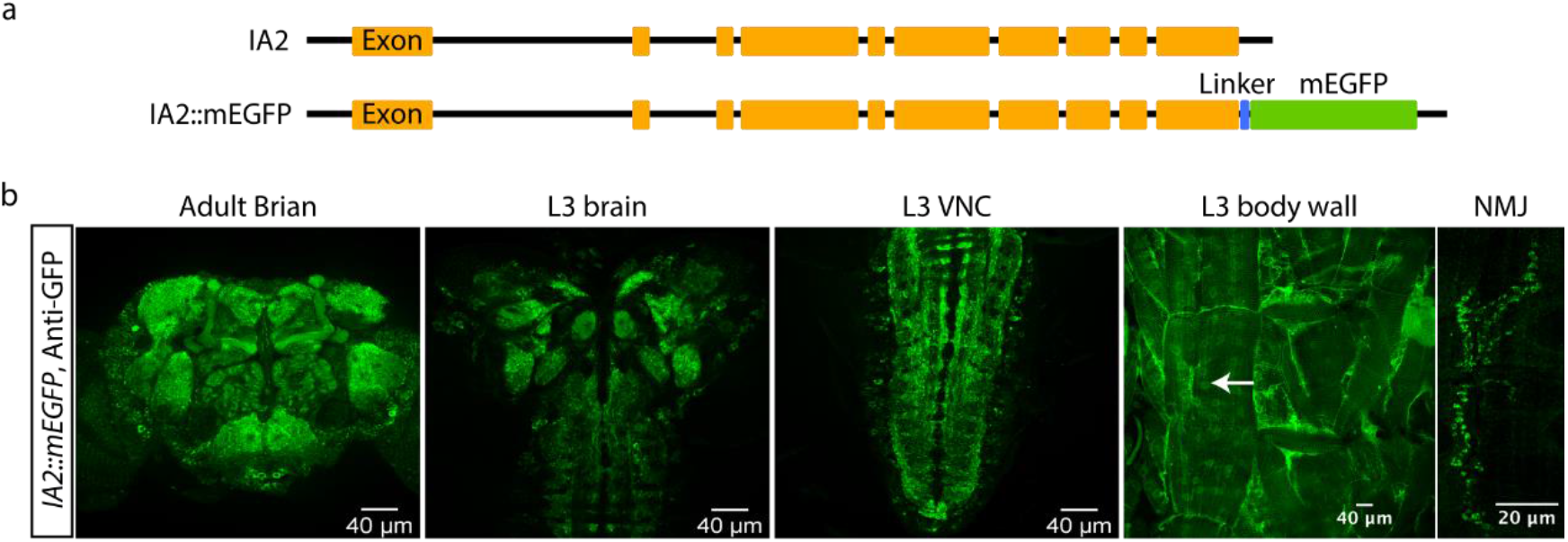
Endogenous IA2 expression patterns in *IA2::mEGFP* fusion animals. **a**, Schematic diagrams illustrate the CRISPR-engineered fusion of mEGFP to the C-terminus of IA2 in the *IA2::mEGFP* fly strain. **b**, GFP staining reveals widespread IA2 expression in the adult brain, third larval instar stage (L3) brain, L3 VNC, L3 body wall and NMJ (left to right). Scale bar: 40 µm, except for 20 µm in the NMJ image.

**Extended Data Figure 2:**
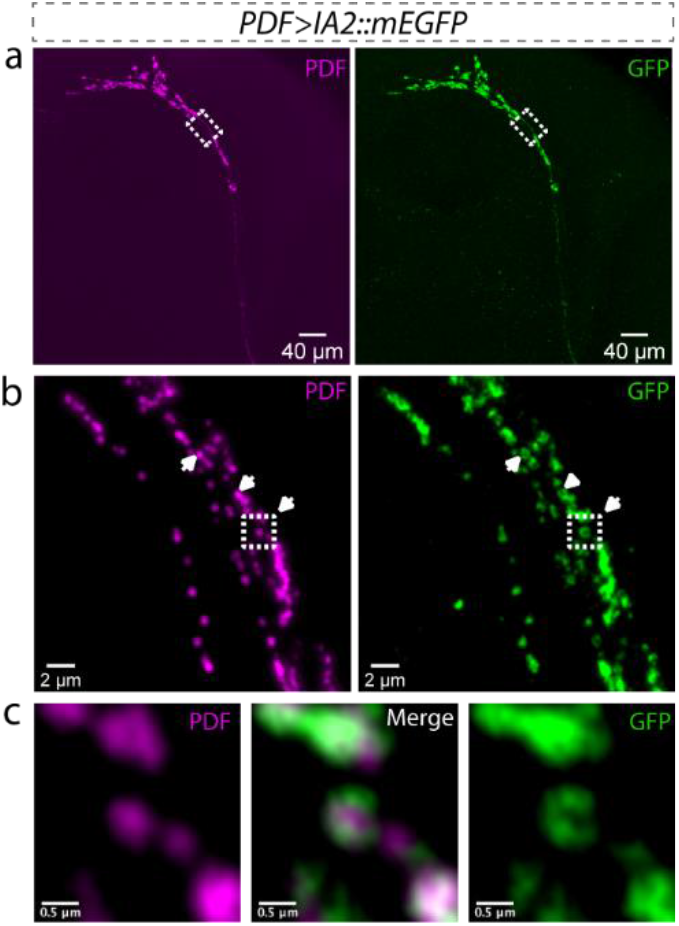
PDF localizes at the center of IA2-labeled DCVs. **a**, Lower magnification images of expanded *Drosophila* brain show the overall profile of PDF-positive neurons (magenta, PDF staining), in which IA2::mEGFP is expressed (green, GFP staining). Scale bar: 40 µm. **b**, Super-resolution images from the outlined area in **a**. Expansion shows that IA2::mEGFP specifically labels the DCVs membrane, with PDF peptide located at the center of the DCVs. Arrowheads indicate examples of single DCVs. Scale bar: 2 µm. **c**, Enlarged images of the outlined area in **b**. Scale bar: 0.5 µm. In **a-c**, magenta indicates PDF and green indicates GFP.

**Extended Data Figure 3:**
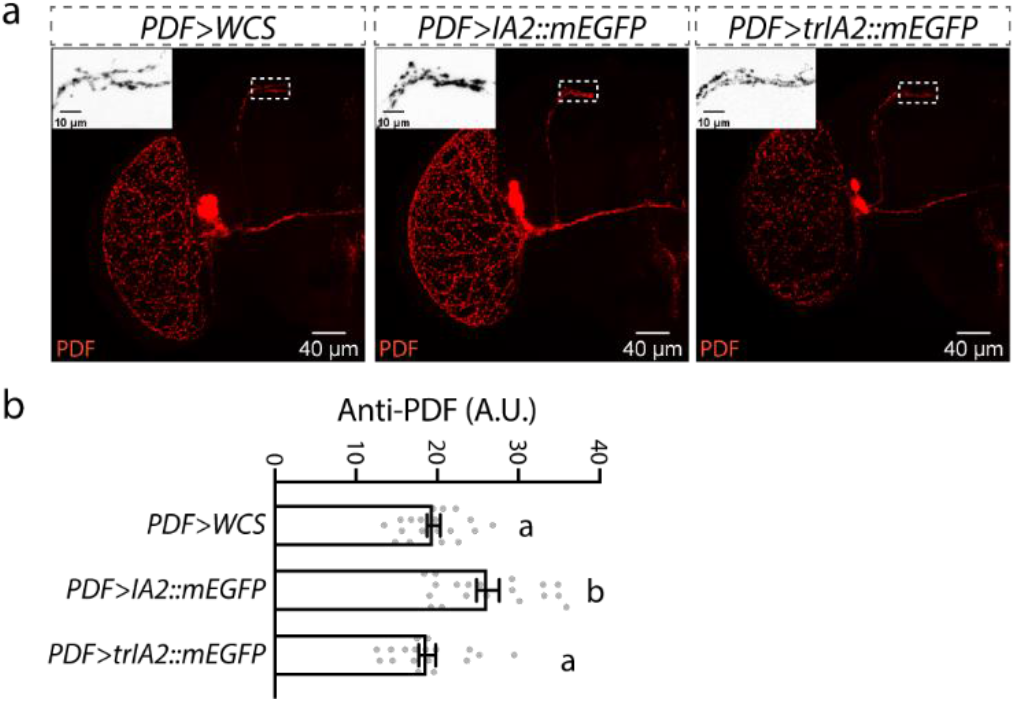
*UAS-IA2::mEGFP* overexpression increases PDF peptide levels in the projections of sLNv neurons, whereas overexpression of *UAS-trIA2::mEGFP* does not. **a**, Representative images of PDF signal in *PDF>IA2::mEGFP, PDF>trIA2::mEGFP* and control *PDF-GAL4* fly brains. Close-up views of the dotted-line outlined regions are shown in the upper left of each image. Scale bar: 40 µm in each panel and 10 µm in each close-up image. **b**, Statistical analysis of PDF peptide levels in the outlined regions from **a**. Data are presented as mean ± SEM, and analyzed by one-way ANOVA with Bonferroni post hoc test as appropriate. Gray dots show individual values. Statistical differences are indicated by letters, and genotypes with the same letter are not significantly different.

**Extended Data Figure 4:**
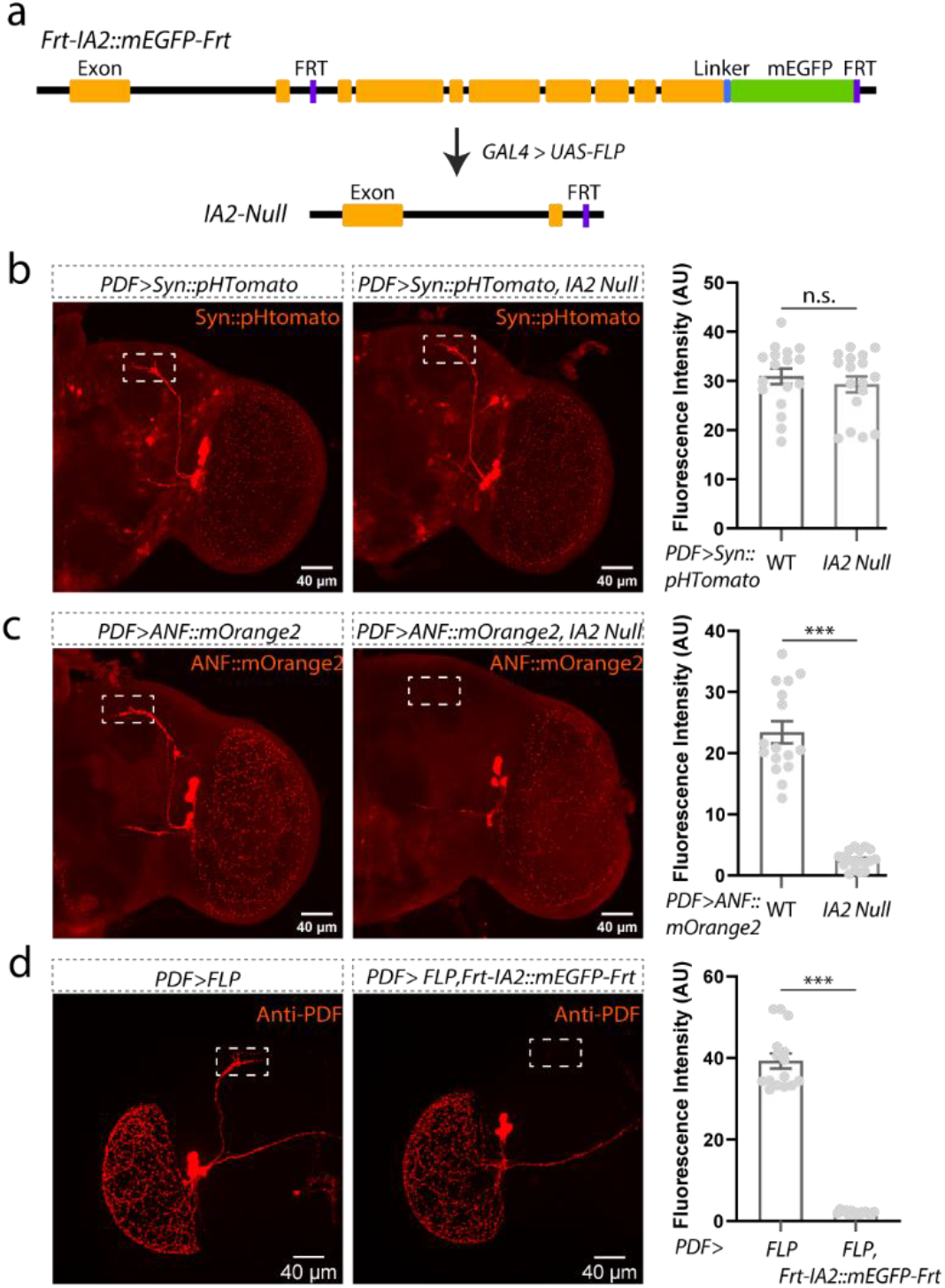
Loss of IA2 does not alter SV levels but does reduce DCVs. **a,** Cartoon of *FRT-IA2::mEGFP-FRT*, a line containing recombination sites that allows deletion of the last 8 exons of the *IA2* gene with expression of flp recombinase. *nos-GAL4* was used to drive recombination and create the null line used in panels **b** and **c**; *PDF-GAL4* was used to delete IA2 specifically in LNvs in panel **d. b,** Representative images from WT and IA2-null brains expressing the SV marker Syn::tdTomato in LNvs. Loss of IA2 does not affect SV numbers. **c,** Representative images of WT and IA2-null brains expressing the DCV cargo ANF::mOrange2 in LNvs. Loss of IA2 significantly reduces DCV number. **d,** Representative images of brains from animals in which IA2 has been removed only from LNvs. Knock out of IA2 in LNvs reduces PDF staining. Scale bar: 40 µm for each panel in **b**-**d**. Data are presented as mean ± SEM, and analyzed by Student’s t test. n.s. indicated no difference; ***P < 0.001. Gray dots show individual values. A.U., arbitrary units.

**Extended Data Figure 5:**
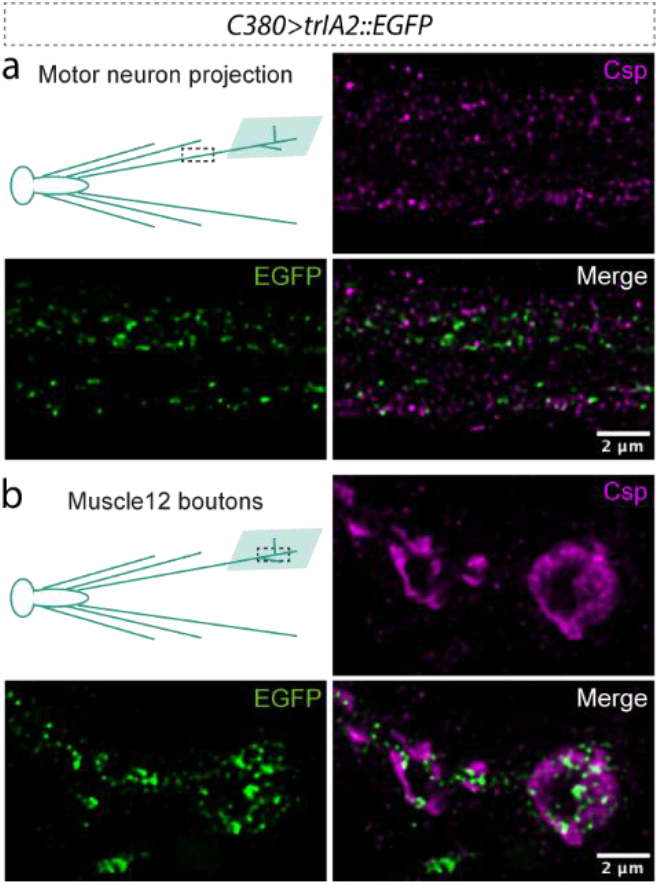
IA2 does not associate with SVs. *C380-GAL4* was used to express *UAS-trIA2::mEGFP* in larval motor neurons. Green indicates trIA2::mEGFP, magenta indicates CSP, an SV marker. **a,** Representative image of a motor neuron axon showing no co-localization of trIA2::mEGFP with CSP. **b,** Representative image of boutons of the CCAP-positive motor neuron 12 showing that trIA2::mEGFP is excluded from regions containing SVs. Scale bar: 2 µm in each panel.

**Extended Data Figure 6:**
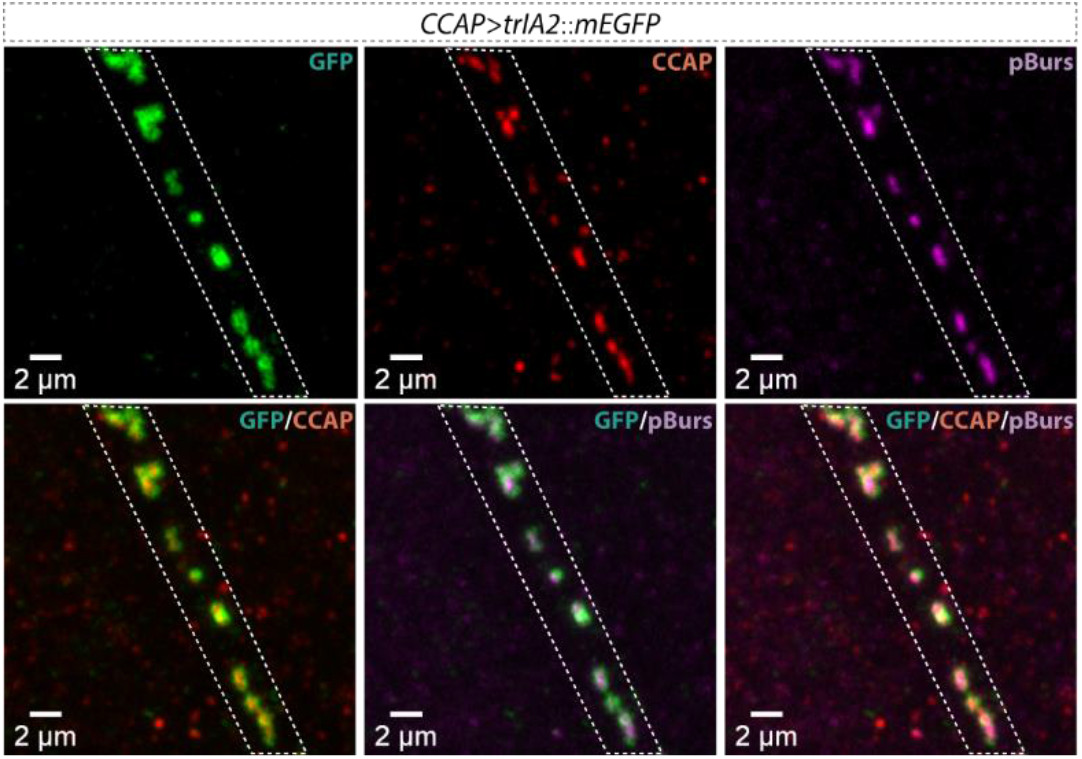
Co-packaging of CCAP and pBurs peptides within individual DCVs in the projections of CCAP-positive motor neurons. The membrane of DCVs is labeled with trIA2::mEGFP. Green indicates trIA2::mEGFP, red indicates CCAP peptide and magenta indicates pBurs peptide. Scale bar: 2 µm in each panel.

**Extended Data Figure 7:**
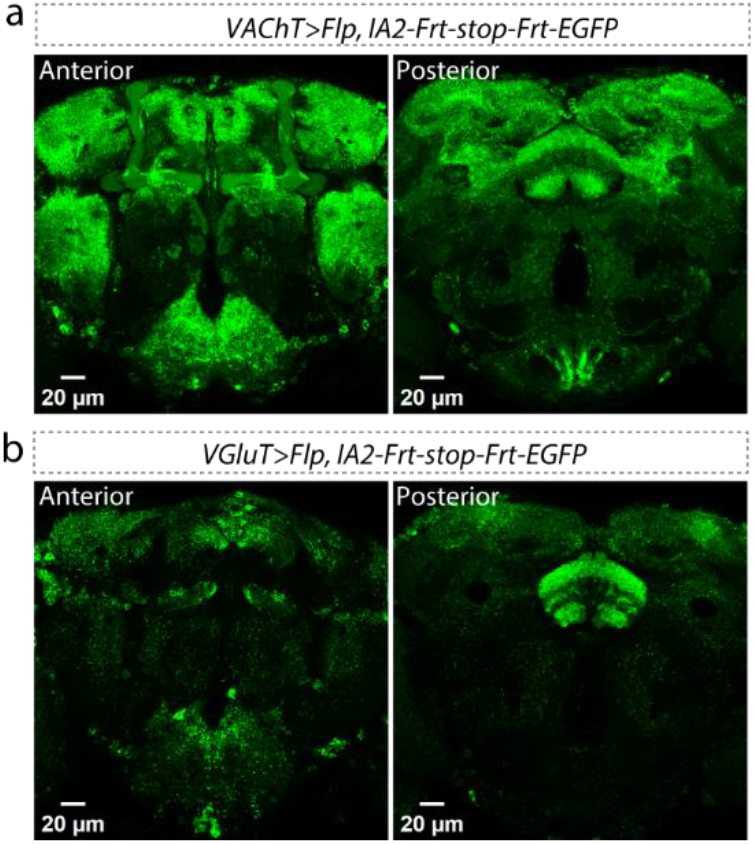
IA2 expression in VAChT-positive (a) and VGluT-positive (b) neurons visualized by conditional tagging of the endogenous gene product. Left panels show the anterior view, and right panels show the posterior view. Scale bar: 20 µm for each panel.

**Extended Data Figure 8:**
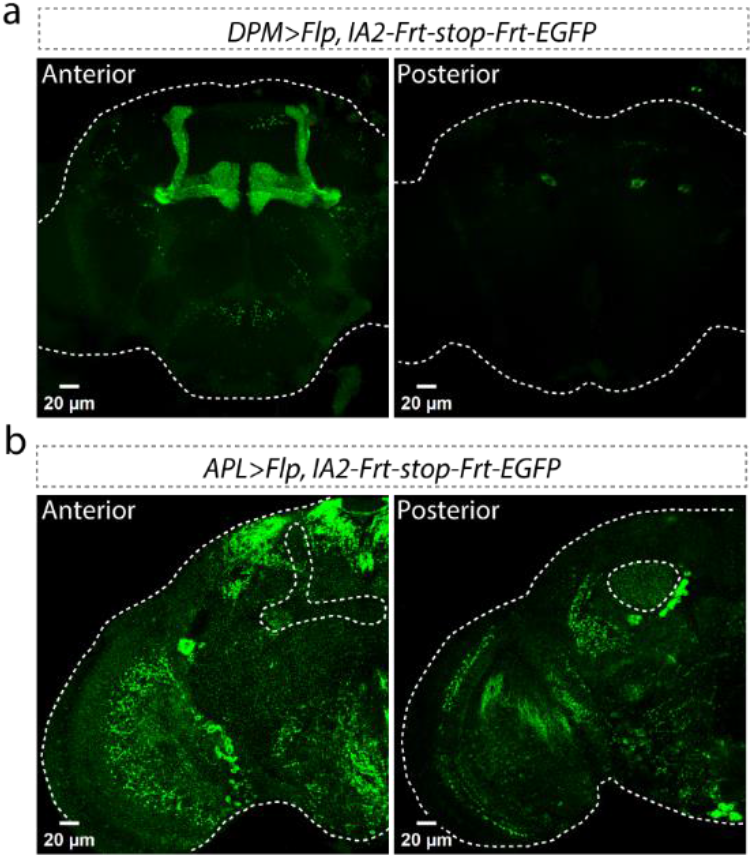
IA2 expression in DPM (a) and APL (b) neurons visualized by conditional tagging of the endogenous gene product. To verify that both DPM and APL neurons normally express IA2, we used conditional tagging. Left panels show the anterior view, and right panels show the posterior view. Scale bar: 20 µm for each panel. Dashed white lines indicate the whole brain in **a**-**b**. The projection of APL neuron in the mushroom body region (left panel of **b**) and the calyx region (right panel of **b**) is outlined by dashed lines.

## References

1 Edwards, R. H. Neurotransmitter release: variations on a theme. Curr Biol 8, R883–885, doi:10.1016/s0960-9822(07)00551-9 (1998).

2 Flavell, S. W., Gogolla, N., Lovett-Barron, M. & Zelikowsky, M. The emergence and influence of internal states. Neuron 110, 2545–2570, doi:10.1016/j.neuron.2022.04.030 (2022).

3 Kim, S., Wallace, M. L., El-Rifai, M., Knudsen, A. R. & Sabatini, B. L. Co-packaging of opposing neurotransmitters in individual synaptic vesicles in the central nervous system. Neuron 110, 1371–1384 e1377, doi:10.1016/j.neuron.2022.01.007 (2022).

4 Woodruff, E. A., 3rd, Broadie, K. & Honegger, H. W. Two peptide transmitters copackaged in a single neurosecretory vesicle. Peptides 29, 2276–2280, doi:10.1016/j.peptides.2008.08.023 (2008).

5 Karagiannis, E. D. & Boyden, E. S. Expansion microscopy: development and neuroscience applications. Curr Opin Neurobiol 50, 56–63, doi:10.1016/j.conb.2017.12.012 (2018).

6 Torii, S. Expression and function of IA-2 family proteins, unique neuroendocrine-specific protein-tyrosine phosphatases. Endocr J 56, 639–648, doi:10.1507/endocrj.k09e-157 (2009).

7 Brand, A. H. & Dormand, E. L. The GAL4 system as a tool for unraveling the mysteries of the Drosophila nervous system. Curr Opin Neurobiol 5, 572–578 (1995).

8 Nishimura, T., Kubosaki, A., Ito, Y. & Notkins, A. L. Disturbances in the secretion of neurotransmitters in IA-2/IA-2beta null mice: changes in behavior, learning and lifespan. Neuroscience 159, 427–437, doi:10.1016/j.neuroscience.2009.01.022 (2009).

9 Johard, H. A. et al. Peptidergic clock neurons in Drosophila: ion transport peptide and short neuropeptide F in subsets of dorsal and ventral lateral neurons. J Comp Neurol 516, 59–73, doi:10.1002/cne.22099 (2009).

10 McDonald, A. J. Functional neuroanatomy of monoaminergic systems in the basolateral nuclear complex of the amygdala: Neuronal targets, receptors, and circuits. J Neurosci Res 101, 1409–1432, doi:10.1002/jnr.25201 (2023).

11 Grygoruk, A. et al. The redistribution of Drosophila vesicular monoamine transporter mutants from synaptic vesicles to large dense-core vesicles impairs amine-dependent behaviors. J Neurosci 34, 6924–6937, doi:10.1523/JNEUROSCI.0694-14.2014 (2014).

12 Lee, P. T. et al. Serotonin-mushroom body circuit modulating the formation of anesthesia-resistant memory in Drosophila. Proc Natl Acad Sci U S A 108, 13794–13799, doi:10.1073/pnas.1019483108 (2011).

13 Harada, K. et al. GABA Signaling and Neuroactive Steroids in Adrenal Medullary Chromaffin Cells. Front Cell Neurosci 10, 100, doi:10.3389/fncel.2016.00100 (2016).

14 Liu, X. & Davis, R. L. The GABAergic anterior paired lateral neuron suppresses and is suppressed by olfactory learning. Nat Neurosci 12, 53–59, doi:10.1038/nn.2235 (2009).

15 Chen, N. et al. Widespread posttranscriptional regulation of cotransmission. Sci Adv 9, eadg9836, doi:10.1126/sciadv.adg9836 (2023).

16 Keles, M. F., Hardcastle, B. J., Stadele, C., Xiao, Q. & Frye, M. A. Inhibitory Interactions and Columnar Inputs to an Object Motion Detector in Drosophila. Cell reports 30, 2115–2124 e2115, doi:10.1016/j.celrep.2020.01.061 (2020).

17 Arslan, A. Extrasynaptic delta-subunit containing GABA(A) receptors. J Integr Neurosci 20, 173–184, doi:10.31083/j.jin.2021.01.284 (2021).

## References

1 Chen, N. Z. Y.; Rivera-Rodriguez E. J.; Yu, A. D.; Hobin, M.; Rosbash, M.; Griffith, L. C. Widespread posttranscriptional regulation of cotransmission. Sci. Adv eadg9836 (2023).

2 Schindelin, J. et al. Fiji: an open-source platform for biological-image analysis. Nat Methods 9, 676–682, doi:10.1038/nmeth.2019 (2012).

3 Chen, F. T. P.W.; Boyden E.S. Expansion microscopy. Science 347, 7 (2015).

4 Inoshita, T., Hattori, N. & Imai, Y. Live Imaging of Axonal Transport in the Motor Neurons of Drosophila Larvae. Bio Protoc 7, e2631, doi:10.21769/BioProtoc.2631 (2017).

